# Mitochondrial phylogeography reveals high diversity and unique divergent lineage in Indian Dugongs (*Dugong dugon*)

**DOI:** 10.1101/2019.12.16.877415

**Authors:** Yellapu Srinivas, Anant Pande, Swapnali Gole, P.V.R. Prem Jothi, K. Madhu Magesh, Sameeha Pathan, Sohini Dudhat, Rukmini Shekar, Devanshi Kukadia, Jeyaraj Antony Johnson, Samrat Mondol, Kuppusamy Sivakumar

## Abstract

1. India plays a central role in dugong conservation by hosting the largest population within south Asia. Current knowledge on status of Indian dugongs is limited due to paucity of reliable ecological data. This study generates mitochondrial control region sequences from about 10% of dugong population from major dugong populations within India. These data was compared with the global data to assess genetic lineages, population structure and genetic diversity of Indian populations.
2. Multiple analyses suggest that the Indian dugong populations are part of a single genetic cluster, comprising south Asia, northwest Indian ocean and southwest Indian ocean populations. Despite small population size, they retain high genetic diversity with unique mitochondrial DNA haplotypes within south Asia.
3. Within India, novel haplotypes were observed from all sampling sites with overall high haplotype diversity (0.85±0.04) but low nucleotide diversity (0.005±0.001). Indian populations exhibit high genetic differentiation with higher within-population variance (63.41%) than among populations (36.59%), signaling population structure. Few haplotypes were shared with Sri Lanka and southeast Asian populations, indicating potential genetic connectivity.
4. Being the most genetically unique population within south Asia, Indian dugong populations are globally significant. We recommend that Indian Dugong populations should be managed as a Conservation Unit to ensure population recovery and long-term survival of the species.

## 1. INTRODUCTION

Dugong (*Dugong dugon* (Müller)) or sea cow is the only strictly marine herbivore species of the order Sirenia. Historically, dugongs were distributed across the tropical and subtropical regions of the Indo-Pacific Ocean, inhabiting shallow coastal waters ranging from east coast of Africa to western Pacific Ocean (Marsh, Penrose, Eros & Hugues, 2002). Currently classified as ‘vulnerable’ by the IUCN Red List, their distribution is restricted to Australia, parts of southeast Asia, Indian subcontinent, Arabian Gulf and eastern coast of Africa (Marsh & Sobtzick, 2015). Within their range, Australian coast retains the largest dugong population, followed by Arabian Gulf. All remaining dugong populations across southeast Asia, Indian sub-continent and east African coast are small and fragmented (Hines et al., 2012; Marsh & Sobtzick, 2015).Their coastal distribution, exclusive dependence on seagrass habitats, long lifespan and slow reproduction rates make them vulnerable to human-mediated impacts across their range. Major threats for dugongs include accidental entanglement in fishing nets, hunting/poaching for meat, vessel strikes, degradation of seagrass habitats and coastal infrastructure development (Marsh, O’Shea & Reynolds III, 2011; Hines et al., 2012; Sivakumar, 2013; D’Souza, Patankar, Arthur, Alcoverro & Kelkar, 2013). Despite various protection measures implemented internationally, their numbers are declining across their range with local extinctions from Mauritius, Maldives and Taiwan (Marsh & Sobtzick, 2015).

India retains the largest dugong population in the south Asia sub-region and thus plays a significant role in dugong conservation at regional and global scale (Sivakumar, 2013). Recent estimates indicate less than 200 individuals distributed in isolated populations along Gulf of Kachchh (Gujarat, west coast), Gulf of Mannar & Palk Bay (Tamil Nadu, south-east coast) and island archipelago of Andaman & Nicobar (Pandey, Tatu & Anand, 2010; Sivakumar, 2013; Sivakumar & Nair, 2013; D’Souza, Patankar, Arthur, Alcoverro & Kelkar 2013). With a decreasing population trend and regionally ‘Endangered’ status, it is absolutely critical to focus on reviving their population and restoring seagrass habitats. However, current ecological knowledge on status of Indian dugongs is limited due to low population densities, rare sighting records and inadequate spatial survey efforts. Lack of long-term monitoring and systematic sampling has also resulted in paucity of reliable information on their genetic lineage in comparison to other dugong populations. So far, genetic status of dugong populations is known from published studies in Australia (Tikel, 1997; McDonald, 1997; Seddon et al., 2014, Blair et al., 2014), Thailand (Palmer, 2004; Bushell, 2013) and western Indian ocean (Plon, Thakur, Parr & Lavery, 2019). These studies indicate three major dugong genetic lineages viz. Australian (restricted and widespread), south-east Asian and western Indian Ocean (Blair, Marsh & Jones, 2013; Plon, Thakur, Parr & Lavery, 2019). Recent work by Plon, Thakur, Parr & Lavery (2019) suggests that the Sri Lankan dugongs are genetically divergent from other populations, based on very limited samples (n=4).Given rapidly declining populations of dugongs in south Asia with recent local extinctions in Mauritius, Maldives and parts of Indian and Sri Lankan coasts (Marsh, Penrose, Eros & Hugues, 2002), adequate sampling to ascertain their position vis-à-vis global populations is of critical importance.

In this paper, we conducted comprehensive genetic sampling of Indian dugongs to describe a) genetic lineage and phylogeography of the Indian dugongs in relation to other dugong populations at global scale and b) the genetic diversity, differentiation and demographic patterns of dugong populations in India. We believe that the results from this study will be critical in developing population specific management plans and thus help in long-term conservation of this globally threatened marine mammal.

## 1. MATERIALS AND METHODS

### 2.1 Permits

The permission to carry out fieldwork and genetic sampling of dugongs was provided by the Ministry of Environment, Forest and Climate Change, Government of India (CAMPA Authority letter number: 13-28(01)/2015-CAMPA). Due to non-destructive sampling approach used in this study, no ethical committee approvals were needed.

### 2.2 Sample collection

Given low population size of dugongs in the Indian subcontinent (<200 individuals), fragmented distribution along the coastal region and rare live sightings (Sivakumar, 2013), it was logistically difficult to conduct systematic sampling of biological material. Hence, most of the sampling was opportunistically conducted from dead stranded dugongs from coastal areas of Gulf of Mannar, Palk Bay, Gulf of Kachchh and Andaman & Nicobar Islands. All samples were stored in ethanol at the field sites and later shipped to Wildlife Institute of India for storage at −20°C until further analysis. In addition to opportunistic sampling, historical samples (bone scrapings) were collected with associated geo-location information from State Forest Departments’ collections. The details of the samples and their geographical sampling location are provided in Supplementary Table 1.

### 2.3 DNA extraction, marker selection and PCR amplification

Total genomic DNA was isolated from all fresh tissue samples using standard protocols mentioned in DNeasy blood and tissue kit (Qiagen, Germany). However, for poor quality museum samples, a modified protocol was used to extract DNA (Mondol, Brufford & RamaKrishnan, 2013). In brief, all samples were cut into small pieces, washed with EDTA and macerated into tiny fragments. The fragments were then completely digested with 40 µl proteinase K for two days, followed by DNA extraction using DNeasy blood and tissue kit (Qiagen, Germany). For museum samples, extractions were carried out in a space dedicated to low quality DNA samples where no earlier dugong DNA extraction has been conducted. Negative controls were included for every batch of extractions to monitor any possible contamination.

We used a universal mammalian primer (A24-Kocher et al., 1989) and dugong-specific primers A58, A77 and A80 (Tikel, 1997) to amplify the mitochondrial DNA (mtDNA) control region from dugong samples. Post-temperature standardizations of these primers, PCR reactions were performed in 10 µl volume with 5 µl Qiagen multiplex PCR mixture (Qiagen, Germany), 0.2 mg/ml Bovine Serum Albumin (BSA), 0.5µM primer mixture and 2µl (2-40 ng/µl concentration) of DNA. PCR conditions included an initial denaturation of 96°C for 15 minutes, followed by 35 cycles of denaturation (96°C for 30 s), annealing (45°C for 30 s) and extension (72°C for 60 s); followed by final extension (72°C for 10 min). During all amplifications, PCR positive and negative controls were included to monitor any possible contamination.

However, it was challenging to amplify the above-mentioned markers in poor quality/degraded bone samples collected in the study. We designed primers to amplify small amplicon sizes from dugong DNA. All published whole mtDNA sequences were downloaded from GenBank with Accession numbers AY075116.1, AJ421723.1, NC003314.1 and were aligned with the sequences generated from tissue samples collected in this study using MEGA v.6.0 (Tamura, Stecher, Peterson, Filipski & Kumar, 2013). Conserved regions within the sequences were visually identified and primers were designed to amplify <250 bp amplicon size to ensure high amplification success from degraded samples. Post-temperature standardization and validation with tissue samples, bone DNA was amplified in 10 µl reaction mixture with 5 µl Qiagen multiplex PCR buffer, 0.2 mg/ml BSA, 0.5µM primer and 2µl of DNA. PCR conditions included an initial denaturation of 96°C for 15 minutes, followed by 40 cycles of denaturation (96°C for 30 s), annealing (50°C for 60 s) and extension (72°C for 90 s); followed by a final extension (72°C for 10 min). During all amplifications, positive extraction control and PCR negative controls were included to monitor any possible contamination. The amplicon sizes of newly designed primers ranged between 140 - 243 bps (see Supplementary Table 2 and Supplementary Figure 1 for more details on primers).

All amplified products were cleaned using Exonuclease-Shrimp Alkaline Phosphatase (GE Healthcare, USA) mixture and sequenced bi-directionally using ABI 3510xl Genetic Analyzer (Applied Biosystems Inc., USA). Sequences were aligned using MEGA v.6.0 (Tamura, Stecher, Peterson, Filipski & Kumar, 2013) and analyzed for missense or frame-shift mutations, possibly arising from sequencing errors. All sequences were then visually examined, matched against GenBank sequences and deposited in GenBank (Accession numbers: MK986797-MK986817).

### 2.4 Data Analyses

Data analyses were performed independently on two dataset alignments:

A) *Global dugong alignment* of 537 sequences (309 bp) consisting of Indian dugong sequences generated in this study (n=21) and sequences downloaded from GenBank (n=516) from previously published studies (Haile, 2008; Jayasankar et al., 2009; Bushell, 2013; Seddon et al., 2014; Blair et al., 2014; Plon, Thakur, Parr & Lavery, 2019). Only 88 sequences (out of 163) were used from Plon et al. (2019) due to ambiguous site calls. The sequences were arranged into global dugong distribution representing Pacific (Australia=383, Papua New Guinea=5, New Caledonia and Palau=1 each); south-east Asia (Thailand=56, Indonesia=7, Japan=2, Philippines and Malaysia/Sabah=1 each); south Asia (India=24, Sri Lanka =4 and Mauritius=2); north-west Indian Ocean (Djibouti=8, Bahrain and Red sea=6 each, Egypt=5, United Arab Emirates=4 and Sudan=2); and south-west Indian Ocean (Tanzania=7, Madagascar=5, east Africa, Mozambique and Kenya=2 each and Comoros=1) regions.

B) *Indian dugong alignment* consisting of 21 sequences (789 bp) of mtDNA control region sequences generated in this study. These sequences were obtained from samples collected from Gulf of Kachchh (n=5), Gulf of Mannar (n=8), Palk Bay (n=4) and Andaman & Nicobar Islands (n=4).

#### 2.4.1 Genetic lineage and phylogeography

The genetic lineage of Indian dugongs’ vis-à-vis global dugong populations and genetic structure within Indian populations was determined using three different approaches:

a. Genetic structure was estimated using Bayesian Analysis of Population Structure (BAPS) v 6.0. (Corander, Marttinen, Sirén & Tang, 2008) to identify the population clusters within both the *global dugong alignment* and *Indian dugong alignment* datasets. Models using spatial clustering of groups followed by admixture analysis were implemented in the program, with K values set between 1 to 15. Each value of K was then analyzed using 500 iterations and 100 burn-ins for each referenced individual per population.
b. Median joining haplotype networkwas constructed using program PopArt v 1.7 (Leigh & Bryant, 2015) to assess the genetic lineage within the *global dugong alignment* as well as for the *Indian dugong alignment* datasets. Haplotype network calculations were carried for both analyses by assigning equal weights to all the variable sites.
c. Phylogenetic relationship of dugong populations was assessed only for the *global dugong alignment*. The best fit nucleotide substitution and partition schemes for the DNA dataset were selected using the Akaike Information Criterion (AIC; Akaike, 1974) implemented in program jModelTest (Posada, 2008). The best-fit substitution model was found to be HKY+I+G. Phylogenetic analysis implemented in program MrBayes v.3.2 (Ronquist et al., 2012) was conducted using Bayesian inference approach (Yang &Rannala, 1997). The chain length consisted of 21 million generations of Markov Chain Monte Carlo (MCMC) simulations, sampled every 1000 generations, with the first 3 million runs discarded as burn-ins. West Indian manatee, *Trichechus manatus* (Accession number: AY963860.1) was kept as an outgroup in the phylogenetic analysis. Finally, FigTree v.1.4 (http://tree.bio.ed.ac.uk/software/figtree/) was used to view and annotate the consensus phylogenetic tree. A posterior probability value ≥ 0.95 and above was considered for indicating strong relationships (Leache& Reeder, 2002).

#### 2.4.2 Genetic differentiation among global dugong populations

Pairwise genetic differentiation values (*F_ST_*) were computed in program Arlequin v 3.5 (Excoffier & Lischer, 2010) for *global dugong alignment* between the dugong population groups derived from multiple structure analysis. Statistical significance was considered with *p*-values < 0.05 between all the pairs of populations after Bonferroni correction for multiple tests.

#### 2.4.3 Genetic diversity estimates, differentiation and demography of Indian dugongs

Genetic diversity estimates including number of haplotypes (H), total number of polymorphic sites (S), nucleotide diversity (π) and haplotype diversity (*h*) were calculated for the Indian dugong dataset (n=21, 789 bp sequence) using program Arlequin v 3.5 (Excoffier & Lischer, 2010). Tajima’s D (Tajima, 1989), Fu’s *F*_S_ (Fu, 1997) and *R2* (Ramos-Onsins & Rozas, 2002) statistics and associated significance values were inferred using Arlequin v 3.5 (Excoffier & Lischer, 2010) and DnaSP v 6.12 (Rozas et al., 2017) to test for demographic signatures. Tajima’s D values are in distinguishable from 0, if the populations are experiencing equilibrium state due to selective neutral variations; D= negative, during demographic expansion or mutational selection and; D=positive, during demographic contraction. Similarly, in the case of Fu’s *F*_S_ statistics (Fu, 1997), negative Fu’s *F*_S_ values are observed during demographic expansion or departure from null hypothesis of neutral selection and population equilibrium. Additionally, mismatch distributions for Indian dugong samples were computed using DnaSP v 6.12 (Rozas et al., 2017) to test whether the populations underwent demographic changes in the recent past. Mismatch distributions display right-skewed uni-modal peaks for populations undergoing expansion whereas ragged and multimodal peaks for populations in demographic equilibrium (Roger & Harpending, 1992). The distribution patterns were further assessed by quantifying the raggedness index (r, Harpending, 1994) and sum of squared deviations (SSD), tested for significance with 10,000 simulations using Arlequin v 3.5 (Excoffier & Lischer, 2010).

Pairwise genetic differentiation values (*F_ST_*) were computed in program Arlequin v 3.5 (Excoffier & Lischer, 2010) between the populations derived from Bayesian clustering analysis. Statistical significance was considered with *p*-values <0.05 between all the pairs of populations after Bonferroni correction for multiple tests. A hierarchical analysis of molecular variance (AMOVA) was conducted using program Arlequin v. 3.5 (Excoffier & Lischer, 2010) to understand genetic variation among individuals, within and between populations as discussed earlier.

## 3. RESULTS

### 3.1 Phylogeography and genetic differentiation in dugongs

The Bayesian clustering, phylogeographic and phylogenetic analyses of the global dugong dataset revealed five major genetic clusters corresponding to a) three clusters grouped in the Pacific region (Australia, New Caledonia, Palau and Papua New Guinea); b) one cluster in southeast Asia (Thailand, Philippines, Japan, Malaysia/Sabah and Indonesia) and c) one cluster comprising of south Asia (Mauritius, Sri Lanka and India), northwest Indian Ocean (Red sea, UAE, Egypt, Sudan, Djibouti and Bahrain) and southwest Indian Ocean (East Africa, Tanzania, Madagascar, Mozambique, Comoros and Kenya) (Figure 1 & 2 and Supplementary Figure 2). While reporting our results, we now use the same nomenclature (Pacific, southeast Asia, south Asia, northwest Indian Ocean and southwest Indian Ocean) in this paper. In total, 76 haplotypes were identified from 537 dugong sequences (Supplementary Table 3). Highest numbers of unique haplotypes were identified from Pacific region (n = 39) followed by southeast Asia (n =20), south Asia (n=8), northwest Indian Ocean (n=6) and southwest Indian Ocean (n=2). Only two haplotypes were shared between these regions; one between Pacific, southeast Asia and south Asia and; another between Pacific, south Asia, northwest Indian Ocean and southwest Indian Ocean (Figure 2 and Supplementary Table 3).

**Figure 1:**
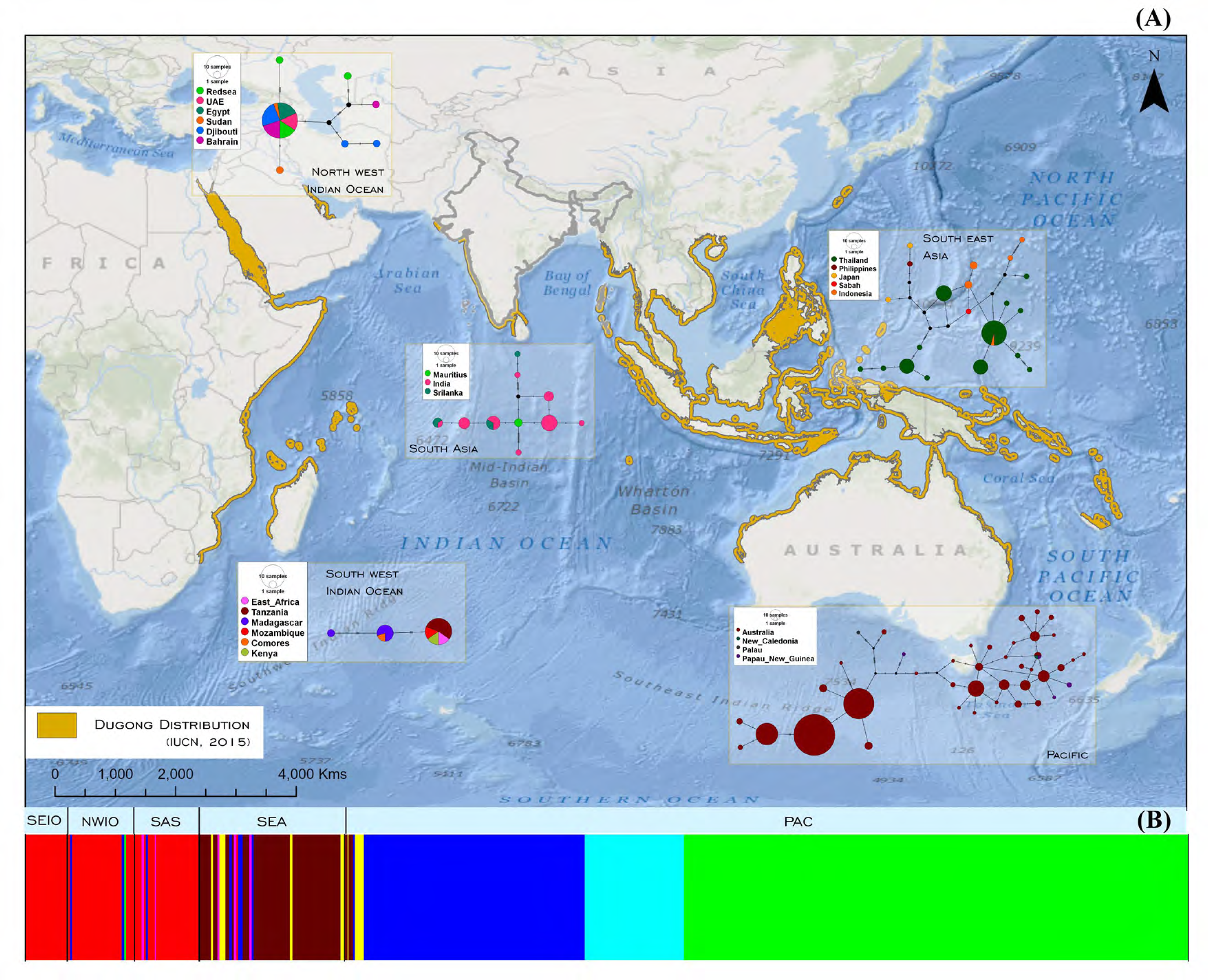
A) Global dugong distribution range and region-wise control region haplotype network generated from dugong sequences. Within each region, circles represent unique haplotypes, different colors represent country of sample origin, and size of the circle represents frequency.B) Population structure using Bayesian clustering (BAPS) analysis for global dugong dataset indicating a total of five clusters using a priori estimate of probable groups. The acronyms are as followed: i) SEIO: southeast Indian ocean; ii) NWIO: northwest Indian ocean; iii) SA: south Asia; iv) SEA: southeast Asia and v) PAC: Pacific regions.

**Figure 2:**
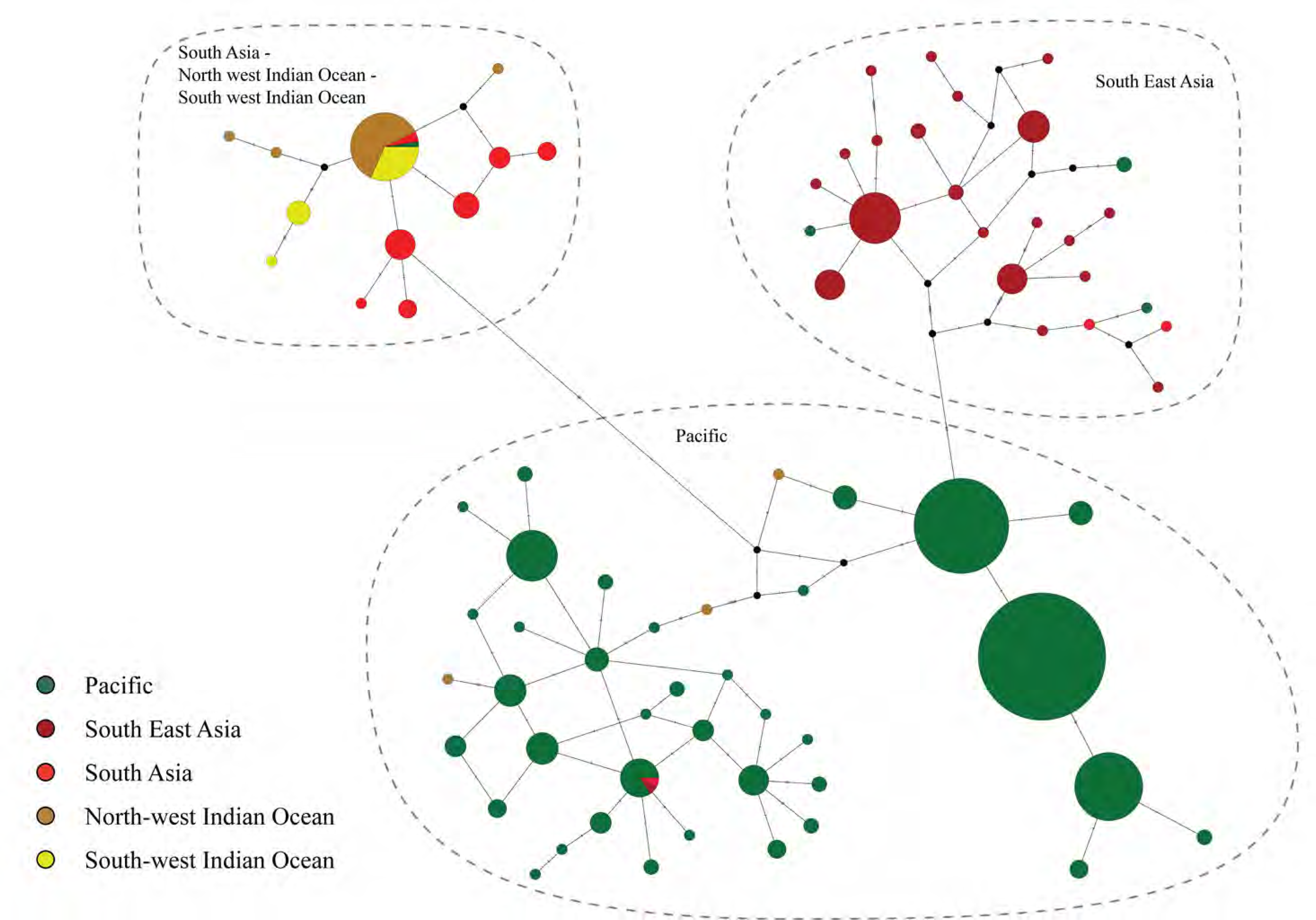
A) Median-joining haplotype network generated from control region sequences obtained from Indian dugong populations. Each circle represents unique haplotype, colors represent sampling locations, and size of the circle represents frequency.Donuts represent sampling locations.B) Donut colors represent proportion of unique haplotypes within each sampling location C) Population structure using Bayesian clustering (BAPS) analysis for Indian dugong dataset indicating a total of three clusters using a priori estimate of probable groups.

The network and phylogenetic analyses indicate a strong phylogeographic structure and divergent mtDNA lineages among different dugong genetic clusters. The genetic differentiation (pairwise *F*_ST_ value) ranged between 0.09-0.66 across different regions (Table 1), indicating variable genetic connectivity among regions. Within each of the dugong distribution regions, relatively high haplotypic diversity (*h* ± SD) was found in south Asia (0.87 ± 0.03), southeast Asia (0.84 ± 0.03) and Pacific (0.81 ± 0.01) in comparison to the southwest Indian ocean (0.49 ± 0.1) and northwest Indian ocean (0.35±0.11) regions. Overall nucleotide diversity (π ±SD) was considerably lower for all regions ranging from 0.01 ± 0.004to 0.02 ±0.01signifying low intra-regional variations (Table 2).

**Table 1:** Pairwise *F*_ST_ values for comparison among global dugong distribution regions

|  | Pacific | South East Asia | South Asia | North West Indian Ocean | South West Indian Ocean |
| --- | --- | --- | --- | --- | --- |
| Pacific | - | 0 | 0 | 0 | 0 |
| South East Asia | 0.25* | - | 0 | 0 | 0 |
| South Asia | 0.42* | 0.60* | - | 0 | 0 |
| North West Indian Ocean | 0.44* | 0.66* | 0.10* | - | 0.01 |
| South West Indian Ocean | 0.43* | 0.64* | 0.16* | 0.09* | - |

Pairwise genetic distance values are mentioned below diagonal while above diagonal are p-values. \*indicates significant differentiation values ( $p < 0.05$ ) after Bonferroni corrections.

**Table 2:** Control region haplotypes and variable sites in Indian dugongs

| Region | Samples<br>(n) | Haplotypes<br>(n) | Polymorphic<br>sites<br>(n) | Haplotype<br>diversity<br>± SD | Nucleotide<br>diversity<br>± SD | *Tajima's D* |  | *Fu's Fs* |  |
| --- | --- | --- | --- | --- | --- | --- | --- | --- | --- |
|  |  |  |  |  |  | value | *p* value | value | *p* value |
| **Pacific** | 390 | 40 | 46 | 0.81 ± 0.01 | 0.02 ± 0.01 | -0.106 | 0.547 | -4.788 | 0.185 |
| **South East Asia** | 67 | 21 | 44 | 0.84 ± 0.03 | 0.02 ± 0.01 | -1.077 | 0.129 | -2.635 | 0.217 |
| **South Asia** | 30 | 10 | 25 | 0.87 ± 0.03 | 0.01 ± 0.01 | -1.215 | 0.099 | -0.249 | 0.487 |
| **North West Indian Ocean** | 31 | 7 | 20 | 0.35 ± 0.11 | 0.01 ± 0.004 | -2.085 | 0.004 | -0.408 | 0.448 |
| **South West Indian Ocean** | 19 | 3 | 7 | 0.49 ± 0.1 | 0.01 ± 0.004 | 0.229 | 0.637 | 3.366 | 0.938 |

### 3.2 Genetic lineage, diversity and demography of Indian dugongs

All three different analytical approaches with the global dugong dataset (n=537, 309 bp sequence) confirm that the Indian dugongs belong to the south Asia- northwest India Ocean-southwest Indian Ocean genetic cluster with unique divergent mtDNA haplotypes. We identified a total of seven unique Indian haplotypes within the south Asia region, where two of them were shared with Sri Lanka (Figure 1 and Supplementary Table 3). One haplotype sampled from Andaman and Nicobar Islands was also observed within southeast genetic cluster (Figure 2).

We generated a longer 789 bp sequence for the Indian dugong samples collected in this study (n=21). A total of eight unique haplotypes were identified from these samples (Figure 3 and Supplementary Table 4). Palk bay, Gulf of Kachchh and Andaman & Nicobar Islands samples showed two haplotypes each, whereas Gulf of Mannar showed four haplotypes. One haplotype was shared between Gulf of Mannar, Palk bay and Gulf of Kachchh. Overall, three haplotypes were unique to Gulf of Mannar, two were unique to Andaman & Nicobar Islands and one each for Gulf of Kachchh and Palk bay, respectively (Supplementary Table 4). Mean haplotype diversity (*h*) and nucleotide diversity (π) was 0.85±0.04 and 0.005±0.001, respectively (Table 3). Our Bayesian clustering analyses (BAPS) indicated three genetic signatures in the sampled areas. Gulf of Mannar has two clusters: one shared with Palk Bay and Andamans whereas the other one shared with Gulf of Kachchh (Figure 3). One sample from Andamans was genetically unique.

**Table 3:**
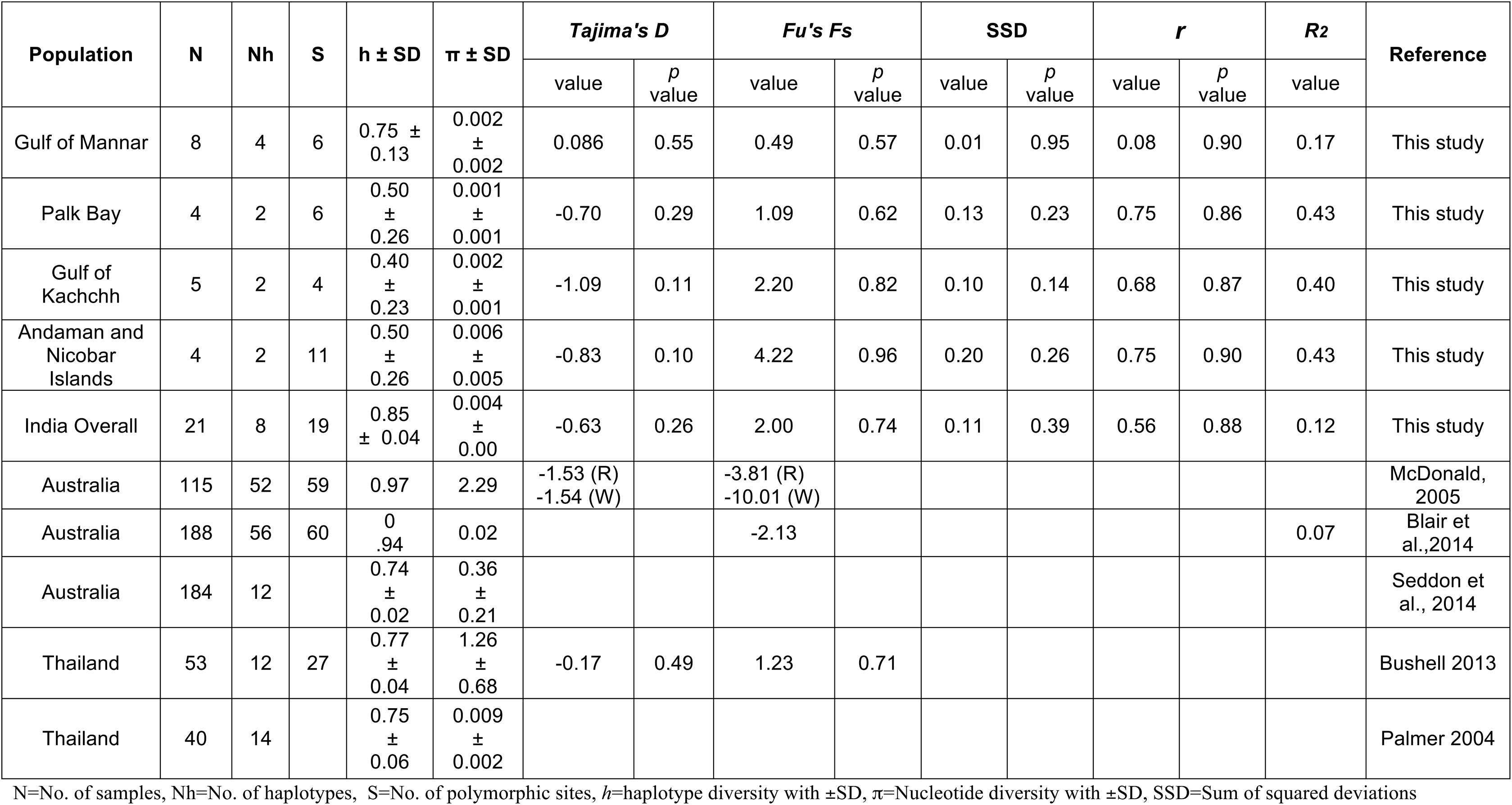
Molecular diversity estimates and demographic changes in Indian Dugong populations

**Supplementary Figure 1:**
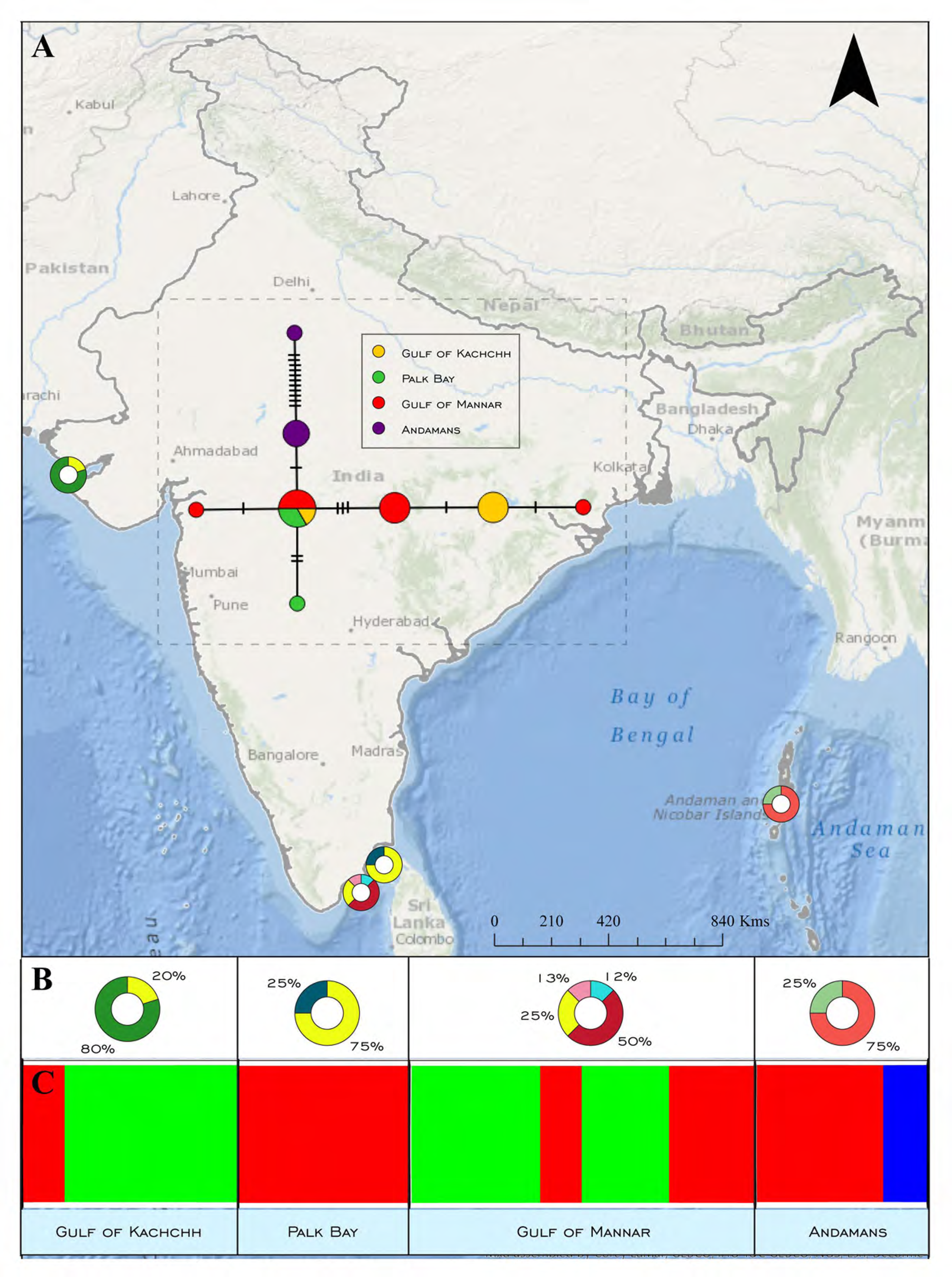
Gel picture showing the lengths of amplification of designed primers with ladder on both the sides

These three areas were genetically differentiated at significant level, with *F_ST_* values ranging between 0.1-0.64 (Table 4). Further, AMOVA results showed higher within-population variance (63.41%) than among populations (36.59%), indicating population structure within the Indian dugongs (see Table 5).

**Table 4:** Pairwise *F*_ST_ values for genetic differentiation between Indian Dugong populations.

|  | **Gulf of Mannar** | **Palk bay** | **Andaman & Nicobar Islands** | **Gulf of Kachchh** |
| --- | --- | --- | --- | --- |
| **Gulf of Mannar** | - | 0.05 | 0.009 | 0.221 |
| **Palk bay** | 0.334* | - | 0.029 | 0.041 |
| **Andaman & Nicobar Islands** | 0.374* | 0.235* | - | 0.014 |
| **Gulf of Kachchh** | 0.103 | 0.639* | 0.487* | - |

Pairwise genetic distance values are mentioned below diagonal while above diagonal are p-values.
\* indicates significant differentiation values ( $p < 0.05$ ) after Bonferroni corrections.

**Table 5:**
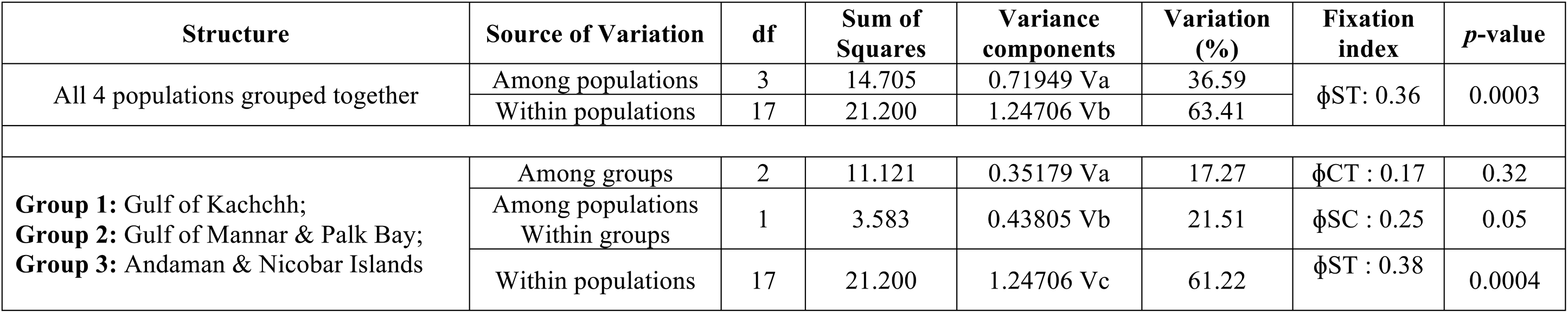
AMOVA values for genetic structuring within and between population groups in Indian dugongs.

Different demography analyses (Tajima’s D, Fu’s F and mismatch distribution) showed contrasting, but non-significant patterns of population demography across these areas. Gulf of Mannar samples showed a positive Tajima’s D value (0.086, p=0.55), indicating population decline, whereas Palk Bay, Gulf of Kachchh and Andaman & Nicobar Islands samples showed negative values (−0.70, p=0.29; −0.83, p=0.10 and −1.09, p=0.11, respectively), indicating population expansion or selection. For Fu’s F, all these regions showed positive values (Gulf of Mannar-0.49, p=0.57; Palk bay-1.09, p=0.62; Gulf of Kachchh-2.2, p=0.82 and Andaman and Nicobar Island-4.22, p=0.96, respectively) indicating population decline (Table 3). Mismatch distribution analyses reveal multi-modal peaks under both constant population size and population growth-decline models, indicating past demographic equilibrium (Figure 4). The observed mismatch values showed a statistically non-significant value of Harpending’s raggedness index *r* = 0.56, p=0.88 (SSD=0.11,p=0.39, *R_2_* = 0.12), indicating a population equilibrium (Table 3).

**Supplementary Figure 2:**
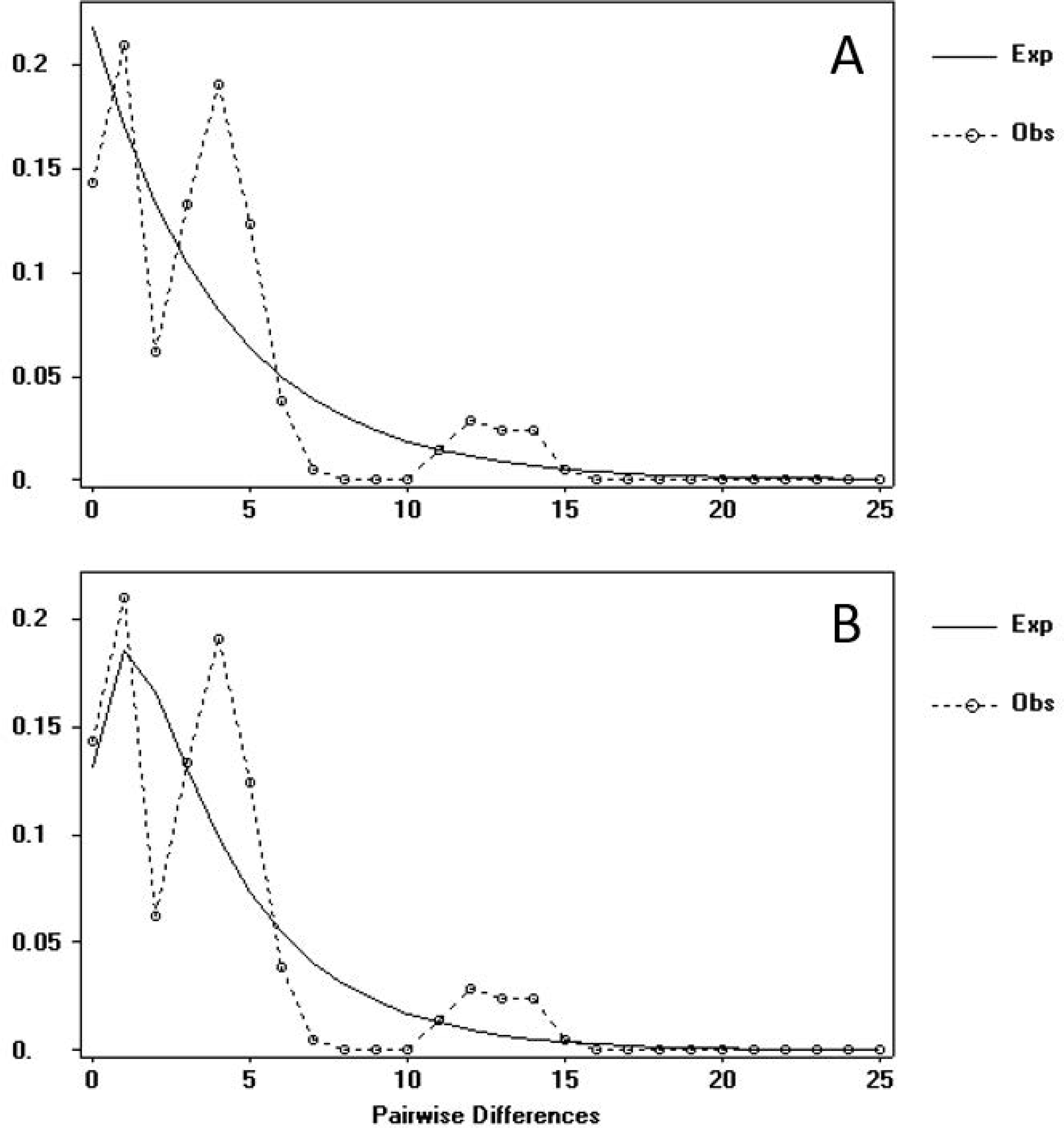
Bayesian phylogenetic tree constructed for global dugong dataset. Values at each node indicates posterior probabilities > 0.50.

## 4. DISCUSSION

This study provides the most exhaustive description of genetic groups of dugong at a global scale till date, and first in-depth investigation of genetic diversity and structure of Indian dugong populations. Given that our sampling represents ∼10% of the available population estimates of dugongs from India (Sivakumar, 2013), the results are crucial for their conservation at regional scale. Earlier work on dugong from Pacific (Tikel, 1997; McDonald, 1997; Seddon et al., 2014, Blair et al., 2014), southeast Asia (Palmer, 2004; Bushell, 2013, Blair et al., 2014) and western Indian ocean (Plon, Thakur, Parr & Lavery, 2019) provided an incomplete picture of genetic groupings due to limited samples from the central part of their global distribution (i.e. south Asia). Our findings fill this gap and show that the Indian samples are part of a single genetic cluster, comprising south Asia, northwest Indian ocean and southwest Indian ocean populations with low genetic differentiation. This pattern of genetic clustering is in accordance with earlier studies by Blair et al. (2013). However, these results show slightly different pattern from Plon et al., (2019), where Madagascar/Comoros formed a unique lineage within the western Indian region. It is also noteworthy to point out that such patterns could also be driven by use of poor quality sequence data from historical samples. We feel that addition of critical dugong samples from India helped in getting a clear picture of genetic groups within this region. Overall, the global data showed a very structured phylogeographic pattern with very limited sharing of haplotypes among the identified regions. While such pattern could arise from incomplete sampling effort across the dugong range, it could also indicate potential loss of gene flow among these regions due to fragmentation of contiguous dugong habitats (Marsh, O’shea & Reynolds III, 2011).

This study elucidates divergent mtDNA lineages of south Asian dugongs within the western Indian ocean populations. Indian dugong population genetically grouped within the south Asia region, although not genetically unique, consists of unique mtDNA haplotypes. Addition of novel mtDNA haplotypes from Indian dugong samples points towards high genetic diversity within south Asia. Two of the previously reported haplotypes from Sri Lanka (Plon, Thakur, Parr & Lavery, 2019) were found to be shared with southern part of dugong distribution in India at Gulf of Mannar, whereas one haplotype sampled from Andaman & Nicobar Islands was observed within southeast Asian lineage. This indicates potential genetic connectivity between these populations in the recent past, and future work should focus on further fine-scale sampling for in-depth investigation. With local extinctions from Mauritius and Maldives and a highly imperiled dugong population in Sri Lanka (Marsh, Penrose, Eros & Hugues, 2002; Marsh & Sobtzick 2015), India holds the largest and potentially the last viable dugong populations within the south-Asia region thereby requiring immediate conservation attention.

Within India, we identified novel haplotypes from each sampling site i.e. Gulf of Mannar, Palk Bay, Gulf of Kachchh and Andaman & Nicobar Islands. Overall haplotype and nucleotide diversity for dugong populations in India were comparable to the Australia (McDonald, 1997; Blair et al., 2014; Seddon et al., 2014) and Thailand (Palmer, 2004; Bushell, 2013) dugong populations. Gulf of Mannar population showed higher haplotype diversity within the Indian regions. Presence of shared haplotypes across Gulf of Mannar, Palk Bay and Gulf of Kachchh suggest potential genetic connectivity among these populations. We found a new haplotype with longer sequences generated in this study (n=789 bp) when compared with earlier studies from Australia (McDonald, 1997; Blair et al., 2014; Seddon et al., 2014) and Thailand (Palmer, 2004; Bushell, 2013), suggesting that longer sequence data is required to assess genetic variation at regional/global scale. Finally, it is important to point out that our Indian dugong data shows contrasting patterns of population differentiation, where we found shared haplotypes among the sampled areas but high *F_ST_* values. Further, the AMOVA analyses indicated signatures of within population structures. We feel that these contrasting patterns probably arise from low sample size from each areas and short sequences (less polymorphic sites) leading to the effects of genetic drift. Further efforts through intensive sampling and more genetic data would clarify these genetic patterns in Indian dugong populations.

Similarly, our demography analyses with mtDNA show contrasting (but non-significant) signals across the sampled area, thus could not be used to deduce population growth or decline. While mitochondrial DNA has been used to assess demographic pattern (for example see Mizuno, Sasaki, Kobayashi, Haneda & Masubuchi, 2018 for Japanese harbor seals), it generally indicates evolutionary signals at longer time frame. Future work should focus on more systematic sampling effort and generate nuclear data (microsatellite, Single nucleotide polymorphisms etc.), which provide much clear signatures of recent population demography (Lah et al., 2016; Komoroske, Jensen, Stewart, Shamblin & Dutton, 2017).

Increasing human population in coastal areas and subsequent coastal developments has adversely affected nearshore ecosystems including seagrass meadows throughout their distribution range (Unsworth & Cullen, 2010), thereby impacting historically exploited seagrass-dependent dugong populations (Hines et al., 2012; Reynolds III & Marshall, 2012). These rapidly declining dugong populations (Marsh, O’Shea & Reynolds III, 2011), are further imperilled by the lack of reliable scientific data on their genetic status. This study addresses the gaps in knowledge on genetic status of Indian dugongs in comparison with the global dugong populations. Indian dugong populations retain global significance being the largest and genetically unique population in the south Asia region (Dugong CMS MoU). Based on these results, we recommend that Indian dugong populations should be managed as a “Conservation or Management Unit” to strengthen conservation strategies for ensuring their long-term survival. With concurrent implementation of Dugong Recovery Program at all the dugong distribution ranges in India (Sivakumar et al., 2018; Sivakumar et al., 2019), we hope that recovering dugongs through a scientifically robust multidisciplinary research and participatory action would safeguard remnant dugong populations in the country.

## 5. ACKNOWLEDGEMENTS

The authors would like to thank the State Forest Departments of Tamil Nadu, Gujarat and Andaman & Nicobar Islands for providing logistics support. We thank the Director, Dean, Research Coordinator and Nodal Officer (External Projects), Wildlife Institute of India for their constant support. A. Madhanraj and Sohom Seal helped in laboratory analysis of samples and GIS analyses, respectively. We acknowledge the generous field support during sampling from the fisherfolk and native communities of the field sites. Authors acknowledge Donna Kwan (CMS Secretariat) and David Blair for their support in initial designing of the genetic work. Funding for this work was provided by the National CAMPA Advisory Council (NCAC), Ministry of Environment, Forest and Climate Change, Government of India (Letter no: 13-28(01)/2015-CAMPA). Samrat Mondol was also supported by the INSPIRE Faculty Award from Department of Science and Technology, Government of India (LSBM-47).

## 8. Author Contribution

KSK, JAJ, SMconceived the study;SG, PVRPJ,KMM,SP,SD,RS, DK collected samples; YS and SM designed the experiments; YS performed the experiments and analyzed the data; YS, AP, SM wrote the manuscript; KSK, SM and AP reviewed the drafts; all coauthorsapproved the final draft.

---

**Supplementary Figure 3:**
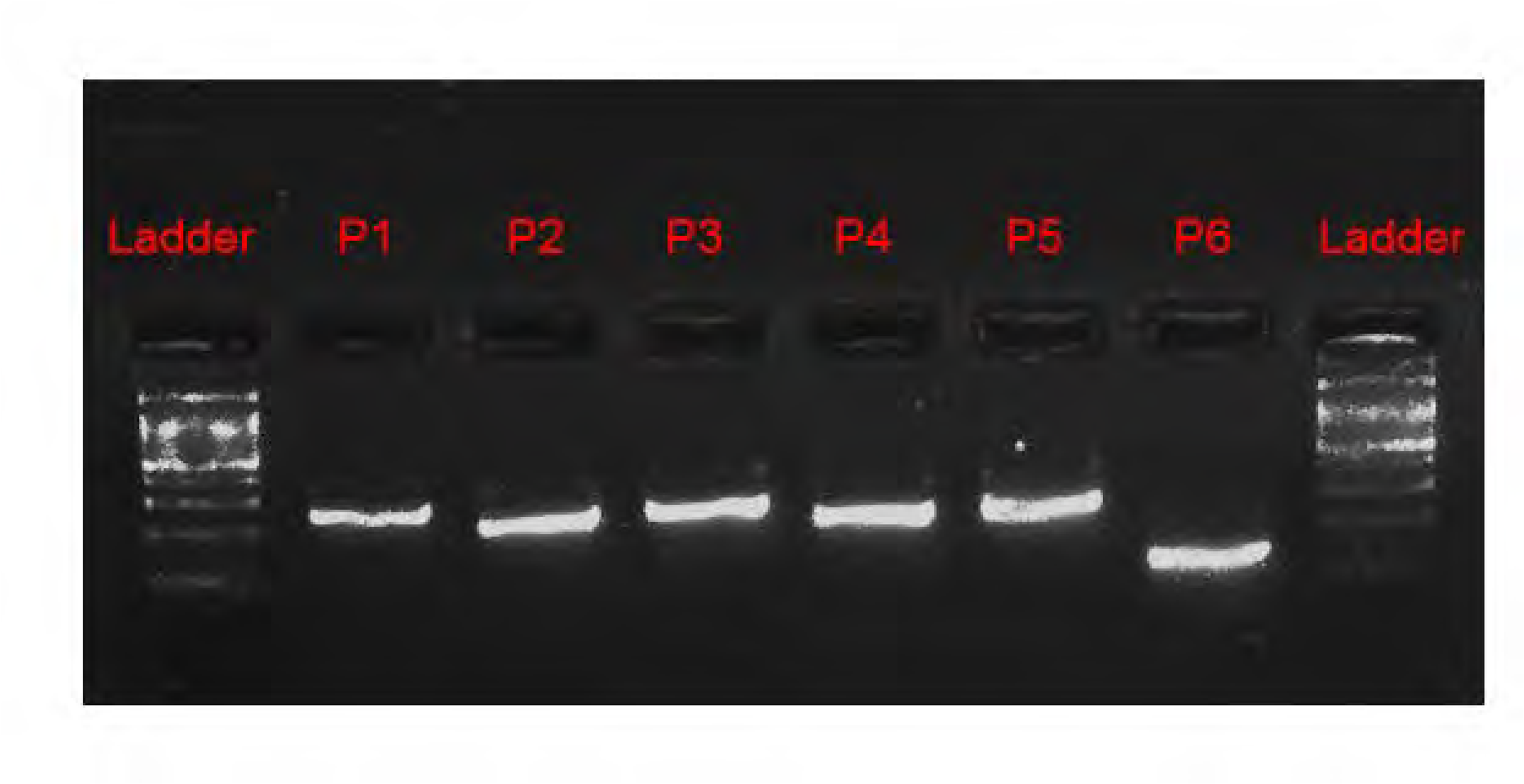
Median-joining haplotype network for global dugong dataset indicating three distinct population groups. Only two haplotypes are shared among these three population groups as depicted in the figure.

**Supplementary Figure 4:**
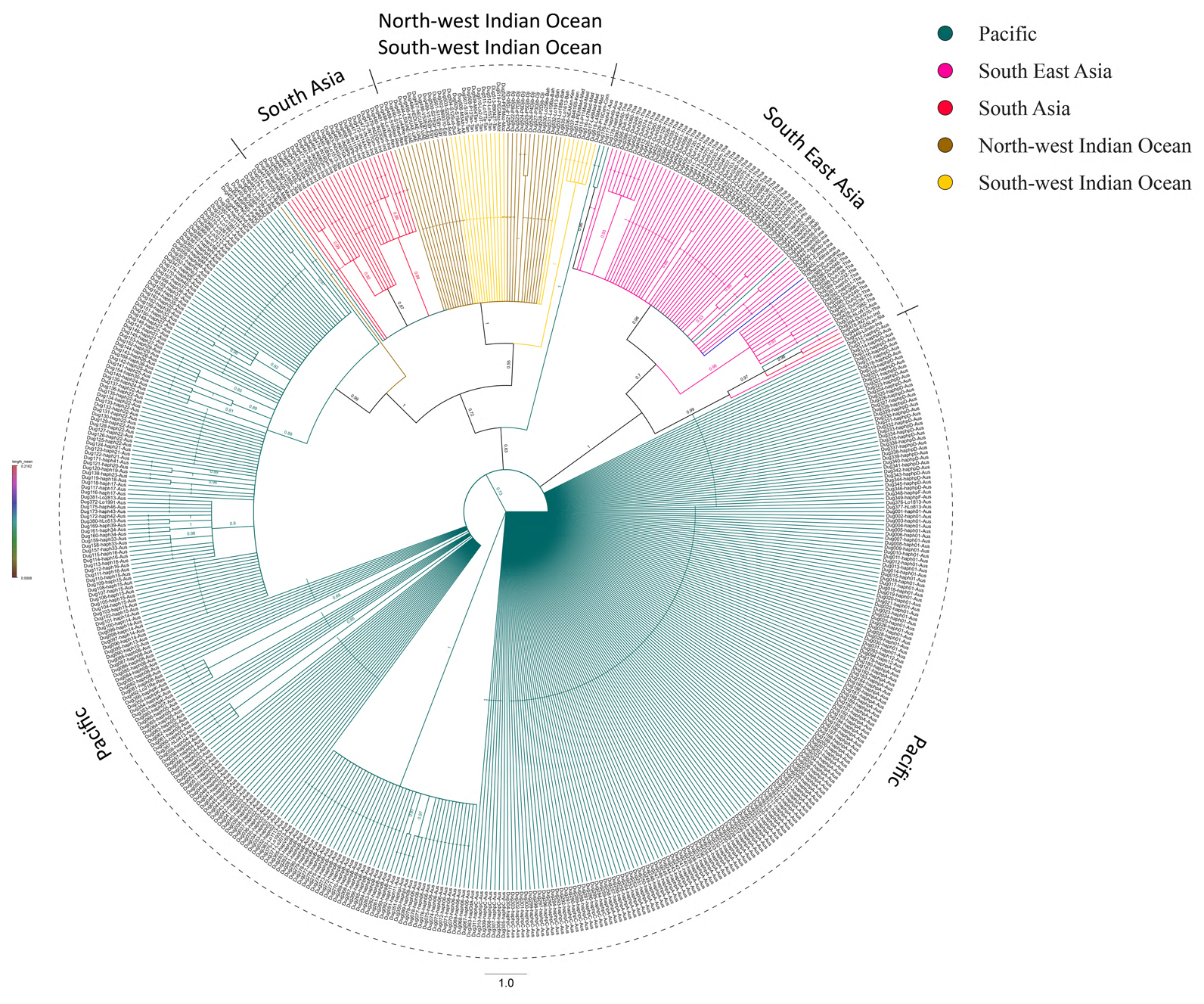
Mismatch distributions of pairwise differences for Indian Dugong populations. Observed (dashed lines) and expected (solid lines) frequencies are depicted using model assuming constant population size (A) and Population growth and decline (B).

**Supplementary Table 1:** Details of samples used in Indian dugong alignment analysis

| **S no.** | **Sample ID** | **Sample Type** | **Sampling Location** | **Year of Sampling** | **GenBank Accession Sequence ID** |
| --- | --- | --- | --- | --- | --- |
| 1 | Dug-1 | Tissue | Gulf of Mannar, Tamil Nadu | 2017 | MK986797 |
| 2 | Dug-2 | Tissue | Gulf of Mannar, Tamil Nadu | 2017 | MK986798 |
| 3 | Dug-3 | Tissue | Gulf of Mannar, Tamil Nadu | 2017 | MK986799 |
| 4 | Dug-4 | Tissue | Gulf of Mannar, Tamil Nadu | 2017 | MK986800 |
| 5 | Dug-6 | Tissue | Gulf of Mannar, Tamil Nadu | 2017 | MK986801 |
| 6 | Dug-7 | Bone | Gulf of Kachchh, Gujarat | 2017 | MK986813 |
| 7 | Dug-8 | Tissue | Gulf of Kachchh, Gujarat | 2017 | MK986814 |
| 8 | Dug-9 | Tissue | Palk Bay, Tamil Nadu | 2017 | MK986806 |
| 9 | Dug-10 | Tissue | Palk Bay, Tamil Nadu | 2017 | MK986807 |
| 10 | Dug-11 | Bone | Palk Bay, Tamil Nadu | 2017 | MK986808 |
| 11 | Dug-12 | Tissue | Gulf of Kachchh, Gujarat | 2018 | MK986815 |
| 12 | Dug-13 | Bone | Port Blair, Andaman & Nicobar Islands | 2018 | MK986809 |
| 13 | Dug-14 | Bone | Port Blair, Andaman & Nicobar Islands | 2018 | MK986810 |
| 14 | Dug-15 | Bone | Port Blair, Andaman & Nicobar Islands | 2018 | MK986811 |
| 15 | Dug-16 | Tissue | Gulf of Mannar, Tamil Nadu | 2018 | MK986802 |
| 16 | Dug-17 | Tissue | Gulf of Mannar, Tamil Nadu | 2018 | MK986803 |
| 17 | Dug-19 | Tissue | Gulf of Kachchh, Gujarat | 2018 | MK986816 |
| 18 | Dug-20 | Tissue | Gulf of Kachchh, Gujarat | 2018 | MK986817 |
| 19 | Dug-21 | Tissue | Shaheed Dweep, Andaman & Nicobar Islands | 2018 | MK986812 |
| 20 | Dug-22 | Tissue | Gulf of Mannar, Tamil Nadu | 2018 | MK986804 |
| 21 | Dug-23 | Tissue | Palk Bay, Tamil Nadu | 2019 | MK986805 |

**Supplementary Table 2:**
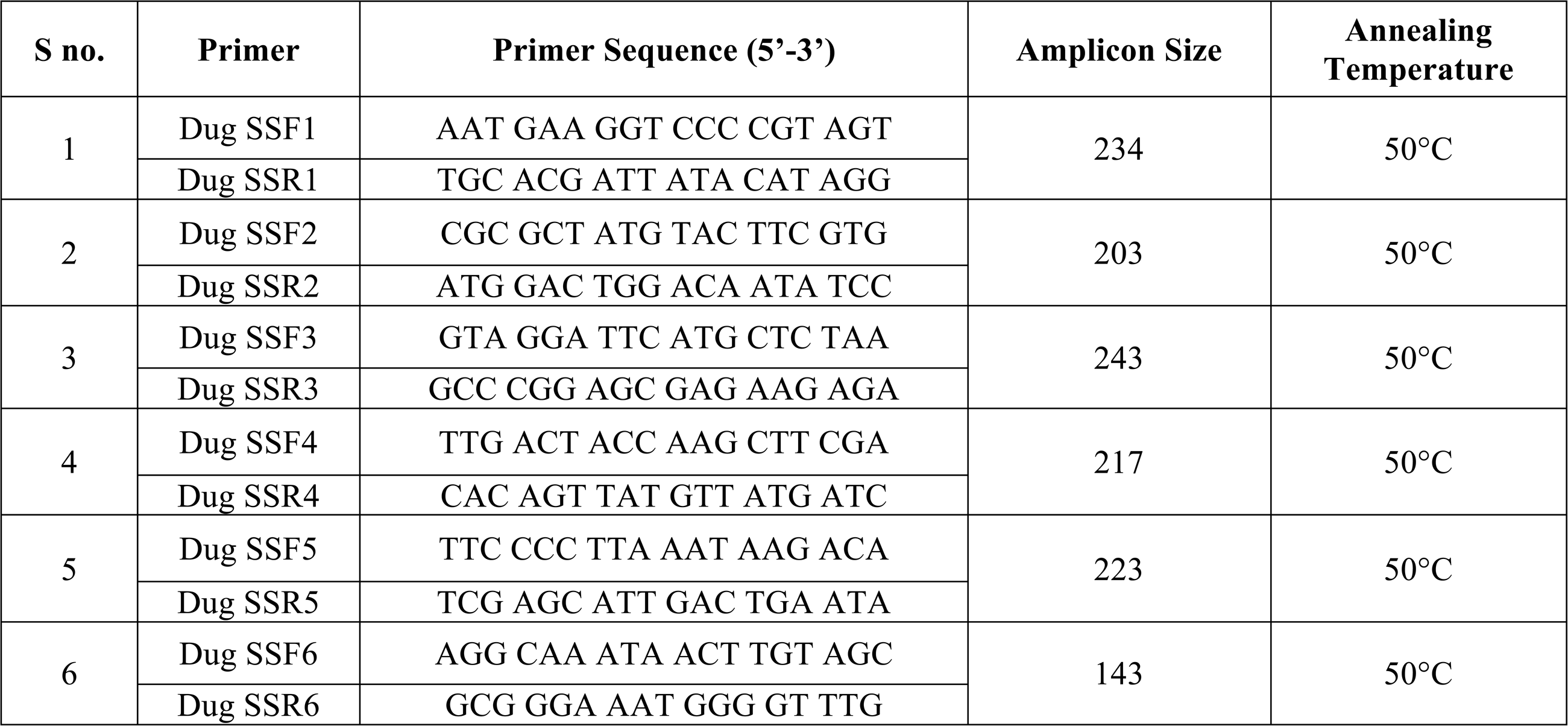
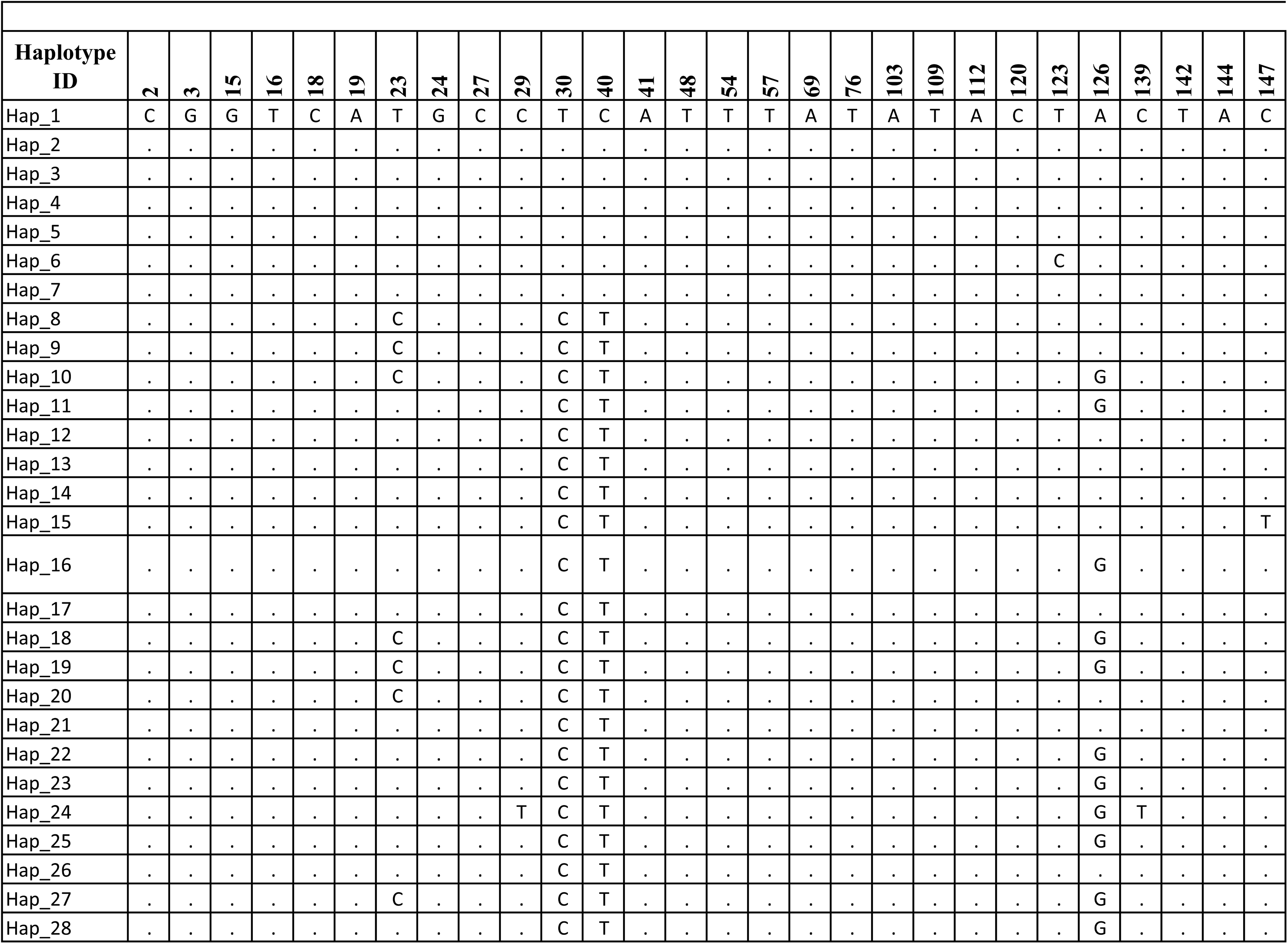
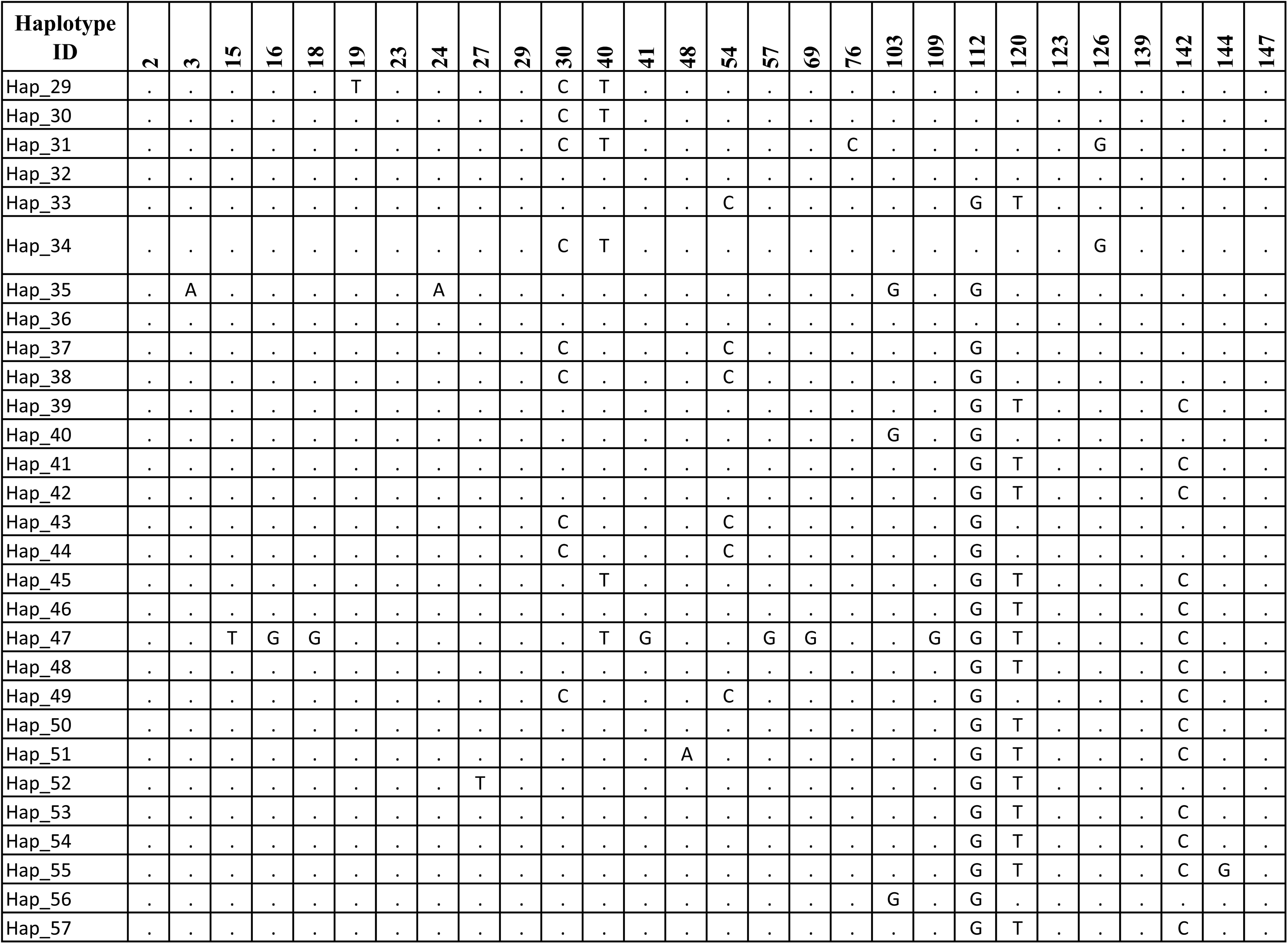
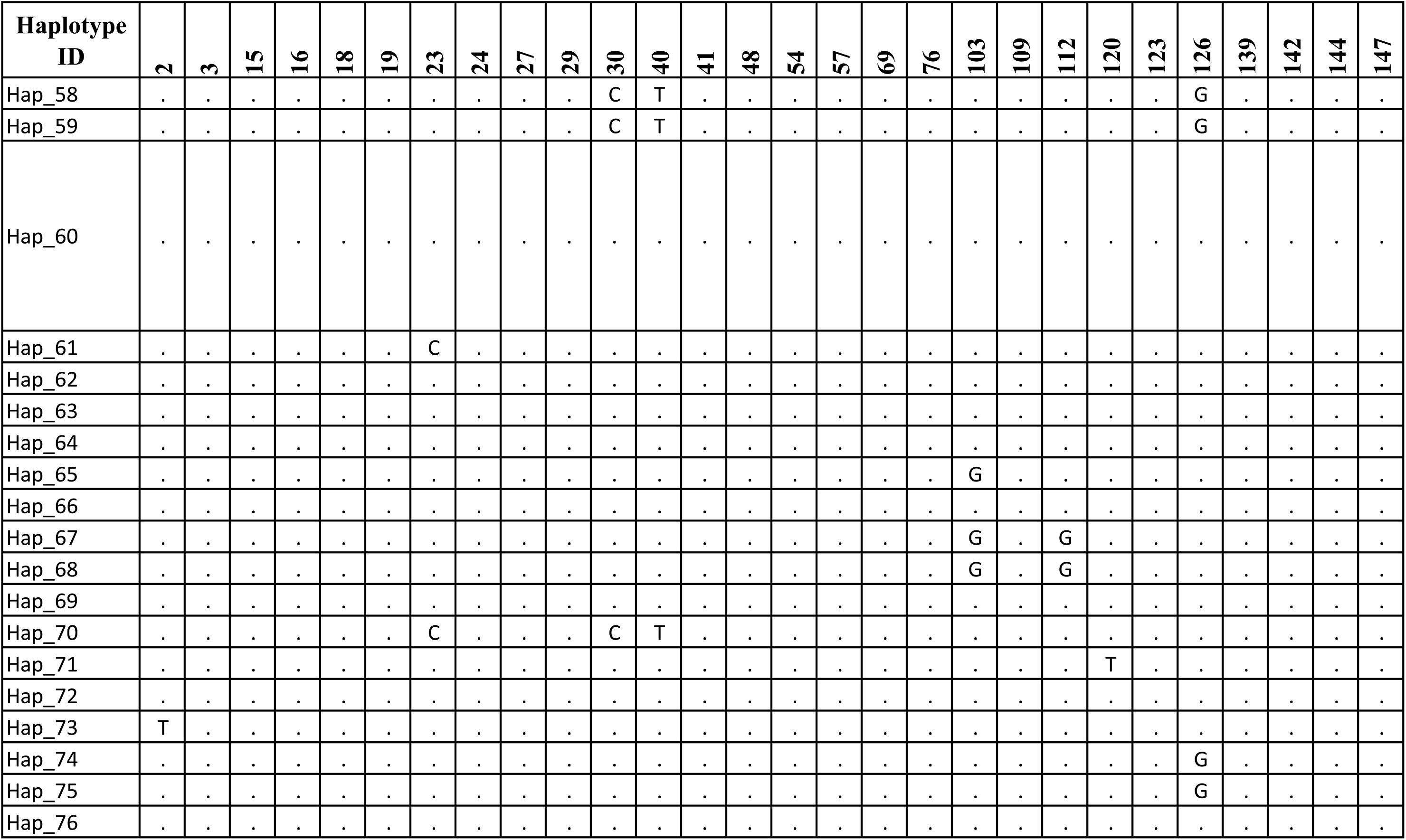
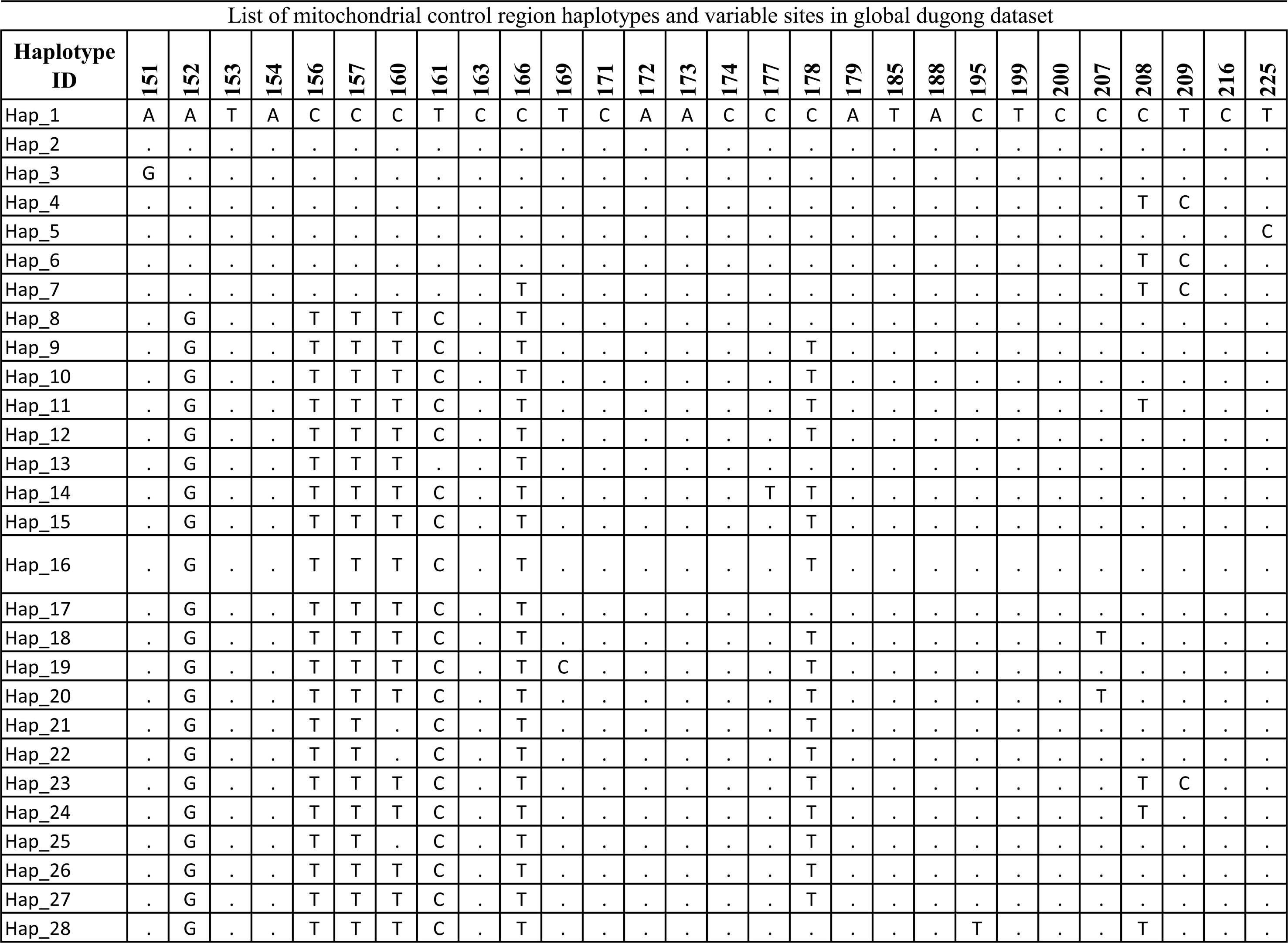
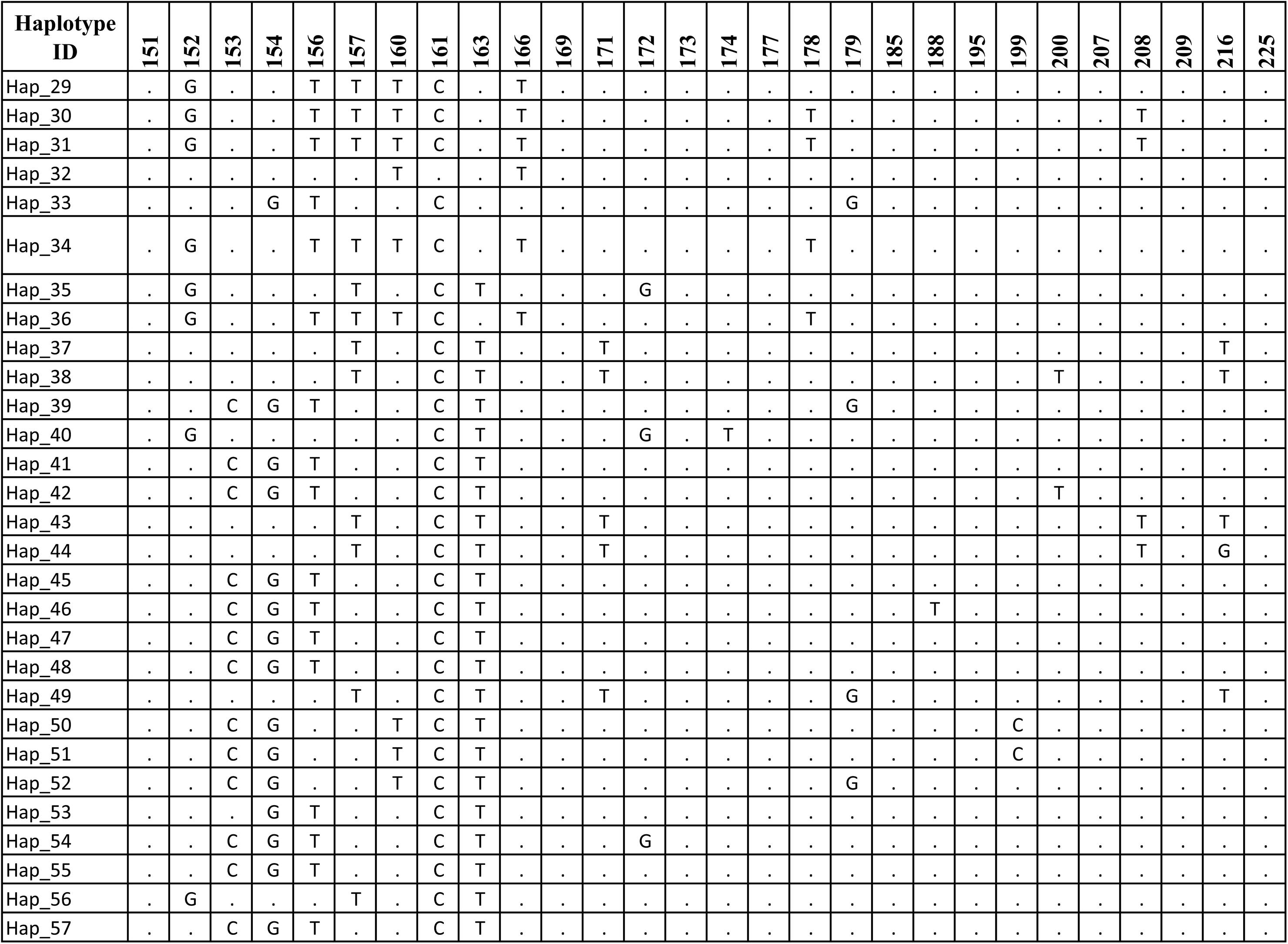
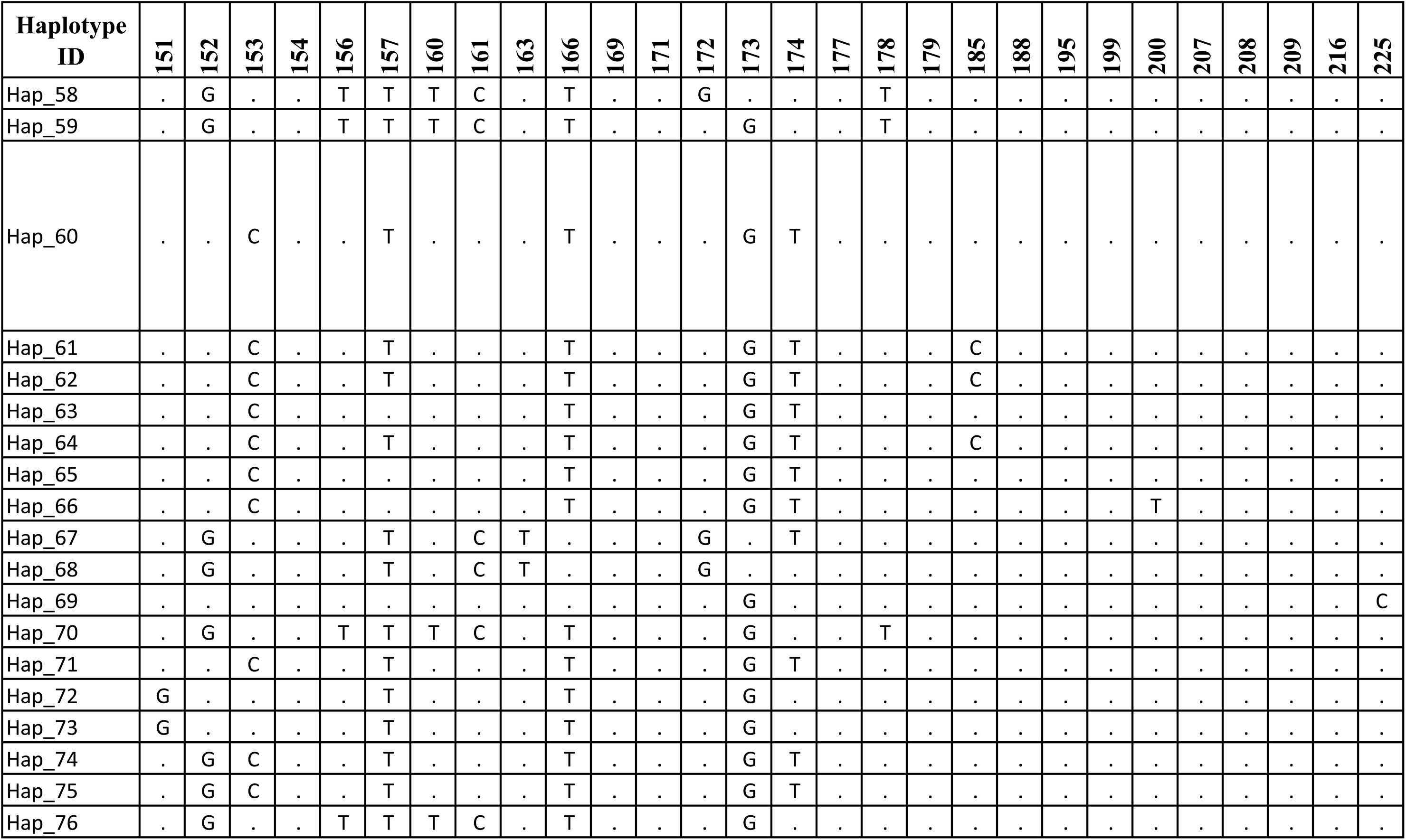
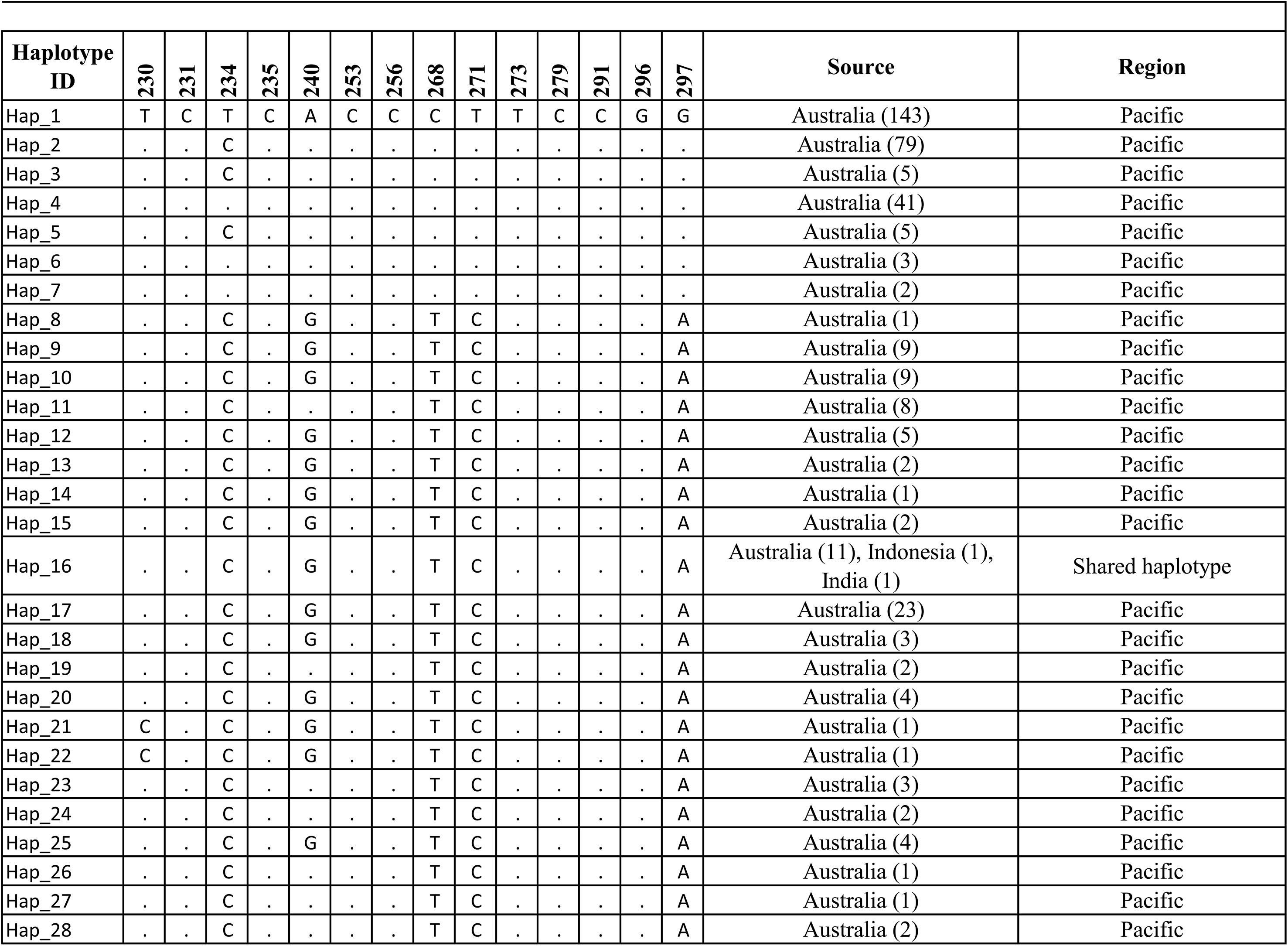
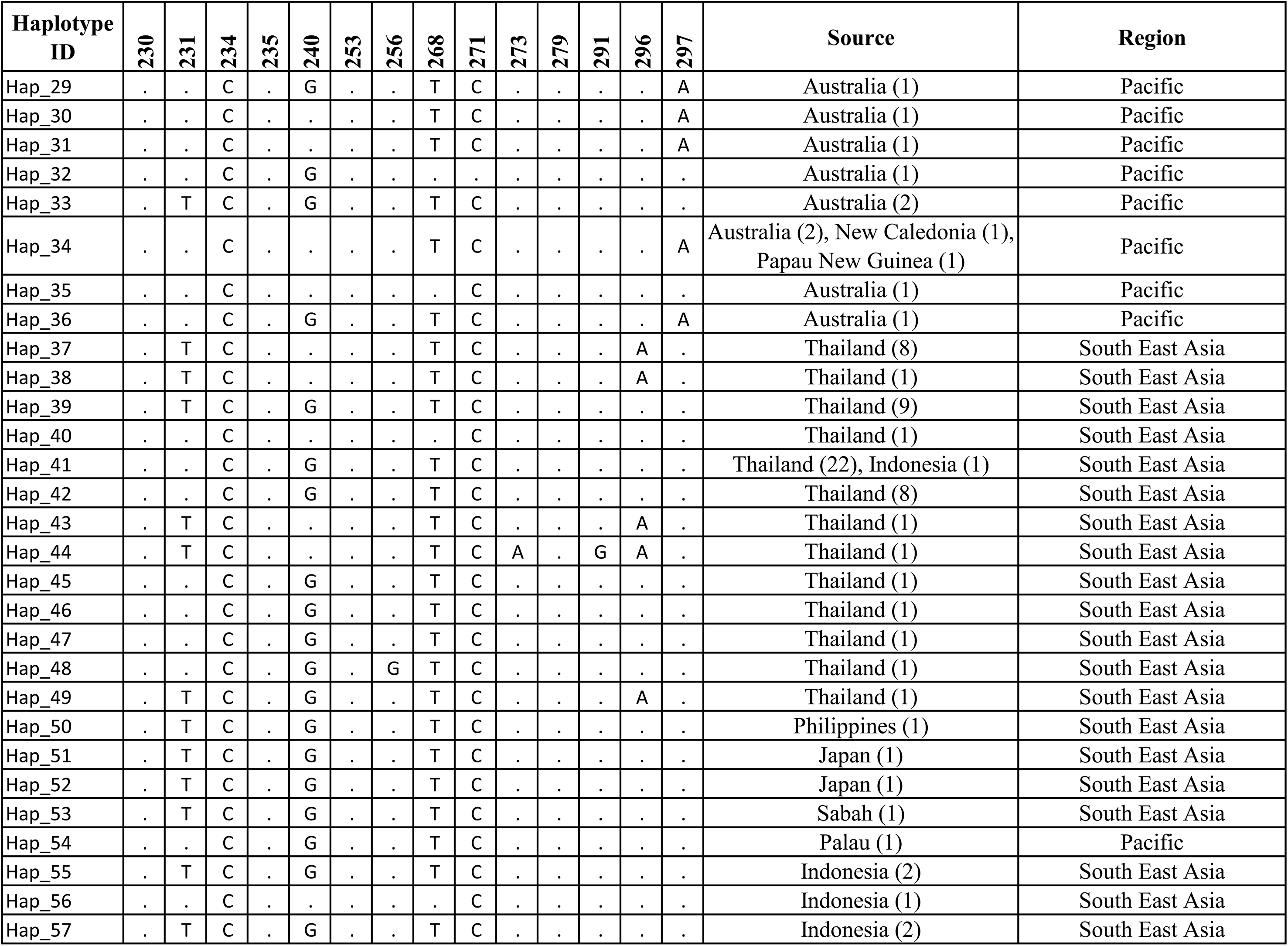
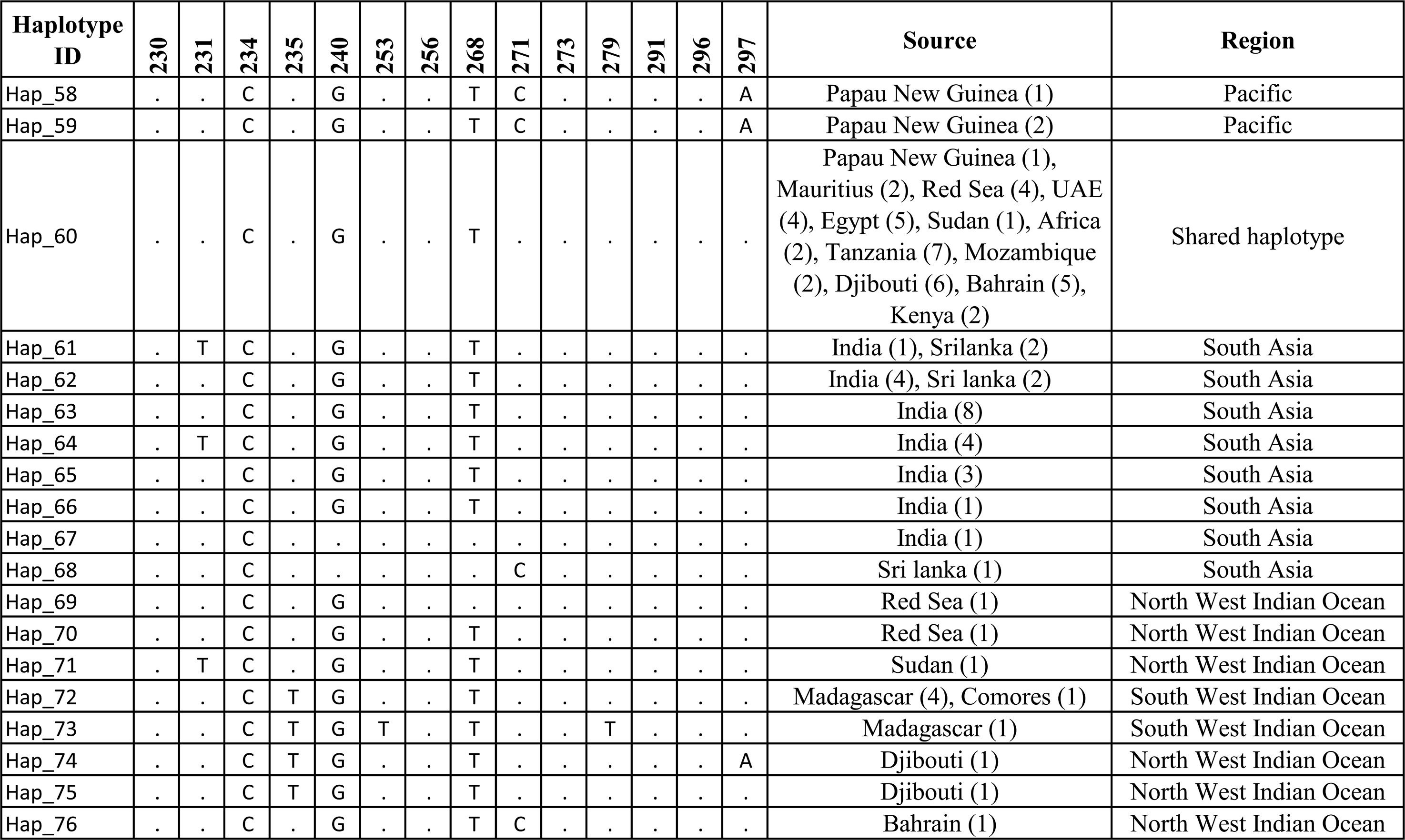
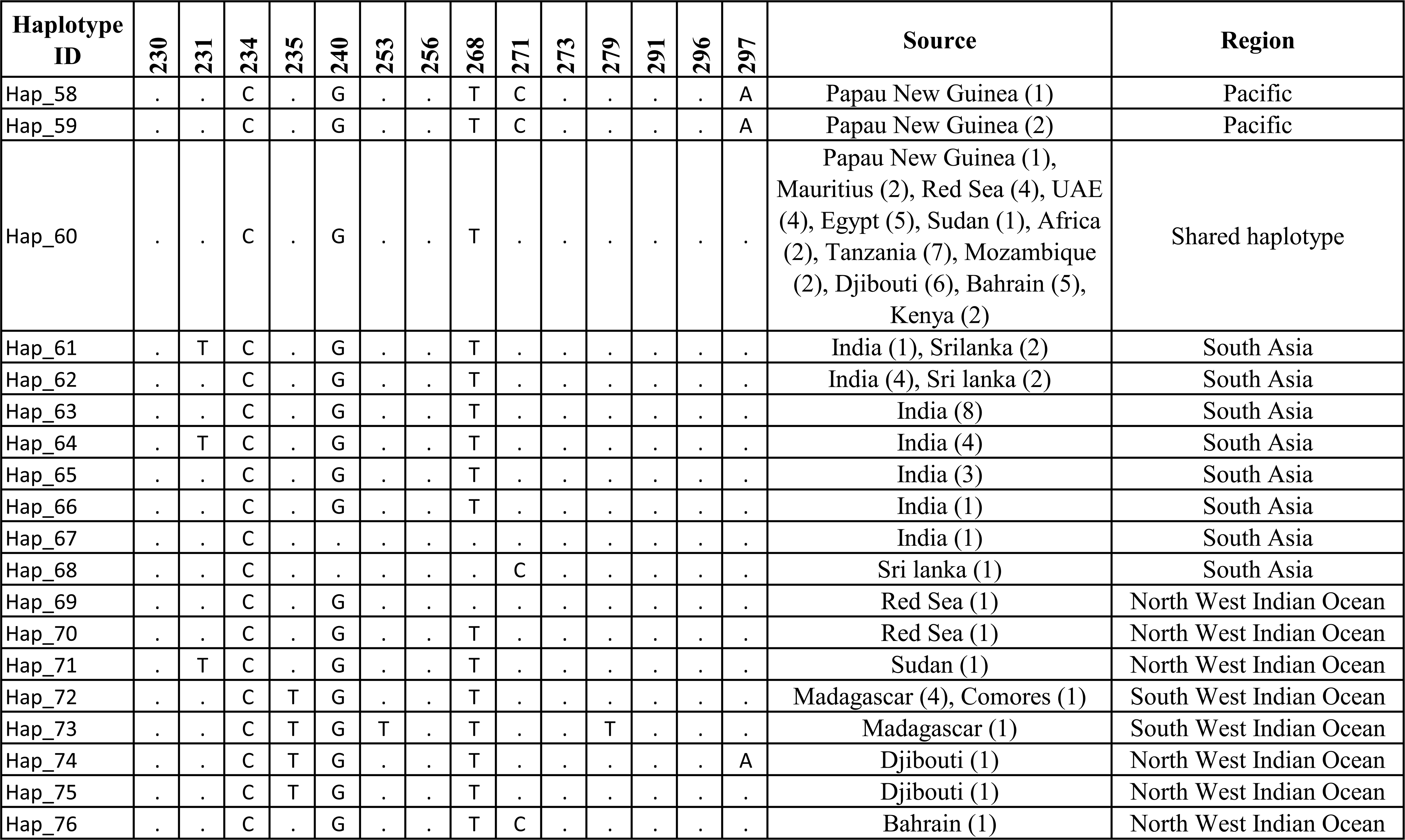
List of primers designed to amplify the mitochondrial DNA control region in dugongs

**Table 2:** Molecular diversity estimates and demographic indices of global dugong distribution regions.

| Haplotype ID | Site numbers |  |  |  |  |  |  |  |  |  |  |  |  |  |  |  |  |  |  | Source | Global ID |
| --- | --- | --- | --- | --- | --- | --- | --- | --- | --- | --- | --- | --- | --- | --- | --- | --- | --- | --- | --- | --- | --- |
|  | 31 | 32 | 136 | 216 | 225 | 265 | 266 | 270 | 274 | 276 | 279 | 285 | 286 | 298 | 313 | 344 | 353 | 381 | 456 |  |  |
| Hap_1 | T | T | C | A | A | A | C | T | T | C | T | A | G | C | C | T | G | T | G | Gulf of Mannar (1) | Hap_61 |
| Hap_2 | . | . | T | . | . | . | . | . | . | . | . | . | . | . | . | C | . | . | . | Gulf of Mannar (4) | Hap_62 |
| Hap_3 | . | . | T | . | . | . | . | C | . | . | . | . | . | T | . | C | . | . | A | Gulf of Mannar (2), Palk Bay (3), Gulf of Kachchh (1) | Hap_63 |
| Hap_4 | . | . | T | . | . | . | . | C | . | . | . | . | . | T | T | C | . | . | A | Gulf of Mannar (1) | Hap_66 |
| Hap_5 | C | C | T | . | . | . | . | C | . | . | . | . | . | T | . | C | . | . | A | Palk Bay (1) | Hap_77 |
| Hap_6 | . | . | T | G | . | . | . | C | . | . | . | . | . | T | . | C | . | . | A | Andaman & Nicobar Islands (3) | Hap_65 |
| Hap_7 | . | . | T | G | G | G | T | . | C | T | C | G | A | T | . | C | A | C | A | Andaman & Nicobar Islands (1) | Hap_67 |
| Hap_8 | . | . | T | . | . | . | . | . | . | . | . | . | . | . | . | . | . | . | . | Gulf of Kachchh (4) | Hap_64 |

Numbers mentioned above each nucleotide base represents positions relative to the common 789 bps sequence used for comparison.

